# Binding to Albumin and Off-Target Toxicity Confound the Use of LRRC8/VRAC Channel Blockers in Cell Physiology Assays

**DOI:** 10.64898/2026.07.13.737859

**Authors:** Mateo A. Boulos, Aayan M. Afghan, Alena Rudkouskaya, Antonio M. Fidaleo, Harmaan S. Sidhu, Meraj T. Khan, Alexander A. Mongin

## Abstract

Volume-regulated anion channels (VRACs), formed by leucine-rich repeat-containing 8 (LRRC8) proteins, are ubiquitously expressed chloride channels essential for cell volume regulation and implicated in diverse physiological and pathological processes. Small-molecule VRAC inhibitors have been reported to modulate paracrine signaling, proliferation, differentiation, migration, and apoptosis, and have been patented for potential therapeutic applications in stroke, cardiovascular and metabolic diseases, and cancer. However, growing evidence indicates that many commonly used VRAC blockers exert substantial off-target effects and frequently fail to reproduce phenotypes observed after deletion of the essential VRAC subunit LRRC8A. Here, we systematically compared effects of several widely used pharmacological VRAC inhibitors with outcomes of molecular downregulation of LRRC8A in limiting proliferation of malignant glioblastoma cells derived from surgical specimens. NIH/3T3 fibroblasts served as a non-malignant control. In serum-containing media, structurally diverse VRAC blockers (DCPIB, DIDS, carbenoxolone, phloretin, and bromadiolone) reduced proliferation in a non-uniform manner, with potencies that did not correlate with reported VRAC affinities and varied markedly among cell lines. Radiotracer-based measurements of VRAC activity indicated that these discrepancies were largely attributable to binding of inhibitors to serum albumin. When experiments were repeated under serum-free conditions, all inhibitors except DIDS and phloretin induced extensive death of both malignant and non-malignant cells, confirmed by microscopy and LDH release assays. This cytotoxicity was accompanied by a marked reduction in intracellular ATP levels, consistent with previously reported mitochondrial uncoupling effects. In contrast, LRRC8A knockdown reduced proliferation without substantial cell death. Together, these findings demonstrate that most commercially available VRAC blockers limit proliferation and viability predominantly through VRAC-independent mechanisms. Under standard culture conditions, serum albumin masks much of their intrinsic cytotoxicity. These results underscore the need for rigorous molecular controls in pharmacological studies and provide basis for developing more selective and less toxic VRAC-targeting agents.

## INTRODUCTION

Volume-regulated anion channels (VRACs) are a family of closely related Cl^−^ channels that are activated by cell swelling and found in virtually all vertebrate cells (Nilius et al., 1997; Okada, 1997; Strange et al., 1996). VRACs are formed by proteins of the leucine-rich repeat-containing family 8 (LRRC8), with LRRC8A (SWELL1) serving as an essential component (Qiu et al., 2014; Voss et al., 2014). Four other homologous proteins, LRRC8B-E, assemble with LRRC8A into heterohexameric channels, with subunit composition determining their diverse biophysical and functional properties (Syeda et al., 2016; Voss et al., 2014). The canonical function of VRACs is emergency cell volume control. In swollen cells, VRAC activation promotes the efflux of Cl⁻ and other inorganic and organic anions and indirectly stimulates K^+^ loss through distinct K^+^ channels, thereby driving the osmotic efflux of water and mediating regulatory volume decrease (RVD) (Hoffmann et al., 2009; Lang et al., 1998; Mongin and Orlov, 2001; O’Neill, 1999). Beyond RVD, VRACs contribute to numerous other aspects of cell physiology, including regulation of membrane potential, cell proliferation, apoptosis, and paracrine signaling (Chen et al., 2019; Eggermont et al., 2001; Jentsch, 2016; Mongin, 2016; Okada, 2024; Thone et al., 2025). Since the discovery of LRRC8, molecular studies have implicated VRACs in glucose sensing and insulin secretion (Kang et al., 2018; Stuhlmann et al., 2018), insulin sensitivity in adipose and skeletal muscle (Kumar et al., 2020; Zhang et al., 2017), spermatogenesis (Luck et al., 2018), immune responses (Concepcion et al., 2022; Lahey et al., 2020; Zhou et al., 2020), and control of brain excitability (Wilson et al., 2021; Yang et al., 2019; Yang et al., 2023). Furthermore, VRAC dysregulation has been linked to several pathological conditions, including stroke, metabolic disorders, and cancer progression (Feustel et al., 2004; Fidaleo et al., 2026; Gunasekar et al., 2022; Kang et al., 2018; Kimelberg et al., 2000; Lu et al., 2019; Zhang et al., 2018; Zhang et al., 2017).

Because of their broad physiological and pathological significance, VRACs have been the subject of sustained pharmacological interest (Figueroa and Denton, 2022; Okada et al., 2019; Pedersen et al., 2016). A variety of broad-spectrum Cl^−^ channel inhibitors, including 5-nitro-2-(3-phenylpropyl-amino)benzoic acid (NPPB), 4,4′-diisothiocyanatostilbene-2,2′-disulfonic acid (DIDS), niflumic acid, and related compounds, inhibit swelling-activated VRAC currents and have been widely used to probe VRAC function in various cells and tissues [reviewed in (Figueroa and Denton, 2022; Okada et al., 2019; Pedersen et al., 2016)]. However, these agents lack specificity, block multiple classes of Cl^−^ channels and produce numerous off-target effects. Efforts to identify more selective tools have yielded several compounds that partially discriminate VRACs from other voltage-gated, Ca^2+^-sensitive, and ligand-gated Cl^−^ channels and/or multidrug resistance transporters. These include the antiestrogens tamoxifen and clomiphene (Maertens et al., 2001; Tominaga et al., 1995; Zhang et al., 1994), the flavonoid phloretin (Fan et al., 2001), the substituted acidic diarylureas NS3623, NS3728, and NS3749 (Helix et al., 2003), and the cysteinyl leukotriene 1 receptor antagonists zafirlukast and pranlukast (Figueroa et al., 2019). Nevertheless, these latter compounds also have multiple targets and exhibit substantial off-target activity. The current best-in-class VRAC inhibitor is the ethacrynic acid derivative 4-[(2-butyl-6,7-dichloro-2-cyclopentyl-2,3-dihydro-1-oxo-1H-inden-5-yl)oxy]butanoic acid (DCPIB) (Decher et al., 2001). DCPIB blocks VRAC currents at 10 µM without affecting voltage-gated, Ca²⁺-activated, or CFTR Cl⁻ channels (Decher et al., 2001). However, subsequent studies revealed that its specificity is also limited, as DCPIB inhibits connexin-43 and pannexin hemichannels, glutamate transporters, suppresses mitochondrial respiration, and either inhibits or activates several classes of K⁺ channels (Afzal et al., 2019; Bowens et al., 2013; Deng et al., 2016; He et al., 2023; Minieri et al., 2013; Zuccolini et al., 2022). Despite these limitations, DCPIB and other less selective VRAC inhibitors remain widely used in physiological studies, often producing conflicting results that complicate interpretation of the role of VRAC in cell physiology.

Arguably, one of the most important but contentious aspects of VRACs physiology is their proposed role in cell proliferation. Numerous early studies reported strong correlations between cell-cycle progression and VRAC current density in both non-malignant and malignant cells and showed that putative VRAC inhibitors—including NPPB, DIDS, NS3728, the antiestrogens tamoxifen and clomiphene, and DCPIB—suppress cell proliferation (Chen et al., 2002; Doroshenko et al., 2001; He et al., 2012; Klausen et al., 2007; Maertens et al., 2001; Shen et al., 2000; Wondergem et al., 2001). Following the molecular identification of VRACs, several groups, including ours, reported elevated LRRC8A expression in multiple human cancers, including colon cancer, esophageal squamous cell carcinoma, hepatocellular carcinoma, gastric cancer, pancreatic adenocarcinoma, and glioblastoma, and demonstrated that LRRC8A deletion reduces several aspects of cancer progression *in vitro* and *in vivo* (Fidaleo et al., 2026; Konishi et al., 2019; Kurashima et al., 2021; Lu et al., 2019; Xu et al., 2022; Zhang et al., 2018). Paradoxically, pharmacological VRAC blockers, including the relatively selective DCPIB, often produces inconsistent or no effects on cell proliferation (Fidaleo et al., 2026; Liu and Stauber, 2019). In studies reporting inhibition of glioblastoma proliferation by DCPIB, the effect was observed only at very high concentrations, 100 µM (Wong et al., 2018). These conflicting observations raise important questions regarding specificity of currently available VRAC inhibitors and their suitability for studying cell proliferation. Here, we sought to address this problem by systematically evaluating several commonly used VRAC blockers—DCPIB, DIDS, carbenoxolone, phloretin, and the recently identified bromadiolone—in longitudinal proliferation assays using patient-derived primary glioblastoma cells. Our findings identify several significant limitations of currently used VRAC inhibitors, underscoring the urgent need for next-generation VRAC modulators with improved specificity, reduced off-target activity, and favorable pharmacokinetic properties.

## MATERIALS AND METHODS

### Materials

Bovine serum albumin (BSA, cat. #A3608), bromadiolone (cat. #46035), carbenoxolone (cat. #C4790), thiazolyl blue tetrazolium bromide (MTT, cat. #M2126), and phloretin (cat. #P7912) were purchased from Millipore-Sigma (St. Louis, MO, USA). DCPIB (cat. #1540) and DIDS (cat. #4523) were acquired from Bio-Techne/Tocris Bioscience (Minneapolis, MN, USA). Lipofectamine RNAiMax transfection agent (cat. #13778) was from Thermo Fisher Scientific (Waltham, MA, USA). All cell culture components, including fetal bovine serum (FBS, cat. #A56707), G5 cell growth supplement (cat. #17503), Dulbecco’s modified Eagle medium (DMEM, cat. #11965), Opti-MEM reduced serum medium (cat. #31985), penicillin/streptomycin (pen-strep, cat. #15140), trypsin-EDTA mix (cat. #25200) and the recombinant protease TrypLE Express (cat. #12605), were of GIBCO brand and acquired from ThermoFisher. All other salts and reagents were purchased from Millipore-Sigma, and were of analytical grade, unless specified otherwise.

### Patient-derived Primary Glioblastoma (GBM) Cell Lines and Fibroblast NIH/3T3 Cells

This study used two primary patient-derived GBM cell lines, here referred to as GBM1 and GBM8, which were isolated and fully characterized in our prior work (Fidaleo et al., 2026; Rubino et al., 2018). In brief, patient-derived GBM cells were isolated from surgical GBM specimens with written patient consent. Following pathological evaluation, leftover GBM tissue fragments were washed in ice-cold sterile phosphate-buffered saline (PBS), transferred into a Petri dish, minced with a scalpel blade, and digested in 0.125% trypsin plus 0.015% EDTA for 10 minutes at 37°C. Cells were separated next by mechanical trituration with a fire-polished glass Pasteur pipette, followed by filtration through a 70-μm cell strainer (Corning). Protease activity was neutralized by adding Opti-MEM supplemented with 10% FBS, and pen-strep. The final cell suspension was then plated in a 75 cm^2^ flask (T75, Corning) and grown in high glucose DMEM supplemented with 10% FBS and pen-strep in an atmosphere of 5% CO_2_/balance air at 37°C. Cell culture medium was replaced twice a week. For the experiments described in this study, GBM cells were utilized between passages 5 and 12.

As an additional non-malignant control, we employed fibroblast NIH/3T3 cells (Jainchill et al., 1969), which were obtained from the American Type Culture Collection (ATTC, cat No. CRL-1658, RRID:CVCL_0594). NIH/3T3 cells were used between passages 2 and 12 of the originally obtained stock. The cultivation conditions and passaging were identical to those described for GBM cells.

### Cell Proliferation Assays

Cell proliferation was assessed using two independent assays. We performed initial screens using a colorimetric MTT assay. All critical findings were independently verified using Coulter counter cell counts.

For the MTT assays, cells were plated in 24-well plates at low densities (20,000 cells/well for GBM1 or 30,000 cells/well for GBM8 and NIH/3T3) and allowed to adhere overnight. Cells were then treated with pharmacological inhibitors as indicated in figure legends. On the second day following initial pharmacological treatment, cell culture media was replaced with fresh media containing the same inhibitors unless stated otherwise. On the third day following pharmacological treatment, cell culture media was removed and replaced with warm Basal medium containing 0.5 mg/ml MTT; cells were then incubated with MTT for 1-2 hours at 37°C. Following the incubation period, MTT-containing media were removed, and the MTT-derived intracellular formazan crystals were then solubilized using dimethyl sulfoxide (DMSO). The optical density of the formazan solutions was determined by measuring the absorbance at 562 nm in an ELx800 plate reader (BioTek Instruments, Winooski, VT, USA). Data are presented as the percentage of MTT conversion relative to untreated control cells on the same plate. All statistical comparisons were performed relative to vehicle-treated (0.1% DMSO) cells on the same plate.

To independently verify the MTT assay results, we directly counted GBM and NIH/3T3 cells using a Coulter counter as described before (Rubino et al., 2018). Briefly, cells grown in 24-well plates at plating density as above were detached from the substrate using the Trypsin and resuspended in the ISOTON II diluent (Beckman Coulter, Miami, FL, USA). Cell numbers were determined using a Z2 series Coulter counter (Beckman Coulter; RRID:SRC_0196761), with the detection cell diameter limits set at 7-17 µm.

### Quantification of cellular ATP content

Intracellular ATP content was determined using a ATPlite detection kit (Revvity, Waltham, MA, Cat. # 6016736) according to the manufacturer’s instructions. GBM1 cells were plated into 96-well plates at the density 10,000 cells per well, allowed to adhere and adapt for 24 h, and then treated with VRAC blockers. After 24-h incubation, cells were lysed using a lysis buffer provided by the manufacturer. The total ATP content in each well was then determined in a luciferase-based reaction in a VICTOR Nivo multimode plate reader (Revvity, RRID: SCR_027779) and experimental luminescence values were compared to an ATP standard curve obtained on the same day.

### Lactate dehydrogenase (LDH) cytotoxicity assay

To assess the cytotoxic effects of VRAC inhibitors, we used the LDH-Glo Cytotoxicity Assay (cat. #J2380, Promega, Madison, WI). Cells were plated in 96-well plates at a density of 8,000 cells per well and cultured for 24 h in serum-free DMEM or OptiMEM supplemented with G5 in the presence of the tested compounds or vehicle. Serum-free medium was used to minimize background LDH signal. After the 24-h incubation period, 5-μL aliquots of extracellular medium were collected to determine LDH activity according to the manufacturer’s instructions. Cells were then lysed with Triton X-100, and 5-μL aliquots of the resulting lysates were processed in the same manner to quantify total LDH. The fraction of LDH released into the medium relative to total LDH was used as a measure of cell cytotoxicity.

### Radiotracer assay of VRAC activity

VRAC activity was quantified by measuring the release of the radiolabeled, VRAC-permeable, non-metabolizable glutamate analog D-[^3^H]aspartate, as previously described (Fidaleo et al., 2026). Briefly, GBM cells were seeded in 12-well plates at a density of 100,000 cells per well. The following day, cells were incubated overnight with 2 μCi/mL D-[2,3-^3^H]aspartate (American Radiolabeled Chemicals, St. Louis, MO; cat. # ARC 212) in DMEM supplemented with 10% FBS at 37 °C in a CO_2_ incubator. Prior to initiation of the release assay, cells were washed three times with serum-containing DMEM to remove extracellular isotope and allowed to equilibrate for 10 min in one of the following media: (a) DMEM + 10% FBS, (b) serum-free DMEM, or (c) DMEM supplemented with 4 mg/mL BSA. Equilibration media have been removed by aspiration and the baseline release was measured by exposing cells to the same control media additionally containing VRAC inhibitors or vehicle. After a 10-min isoosmotic incubation, extracellular media were collected for determination of radiotracer release and replaced with hypoosmotic media to stimulate VRAC activity. Hypoosmotic conditions were achieved by reducing osmolarity by 30% through dilution with water while maintaining the same FBS or BSA concentration. Cells were incubated under hypoosmotic conditions for an additional 10 min, after which the medium was collected for analysis. Radiotracer release was quantified using a TRI-CARB 4910TR liquid scintillation counter (Revvity, RRID:SCR_027779) after mixing collected media with Ecoscint A scintillation fluid (National Diagnostics, cat. #LS-273). D-[^3^H]aspartate release was normalized to the remaining intracellular radiotracer content, which was determined for each well following cell lysis in a solution containing 2% SDS and 8 mM EDTA.

### Statistical Analyses

All data are presented as mean values ± SD, with the number of independent experiments (n) indicated in each figure. In majority of cases, ‘n’ represents an independently prepared and treated cell culture. For cell proliferation and VRAC activity assays, each condition was tested in four different wells, which were averaged to yield a single value (n = 1). In few instances, ‘n’ corresponds to an independently treated or independently transfected well on the same multi-well cell culture plate. Data distribution in experimental datasets was tested for normality. In most cases, statistical differences between normally distributed groups were assessed using one-way ANOVA followed by Dunnett’s post hoc test for multiple comparisons. Comparisons between two groups were performed using an unpaired t-test. In experiments where data were normalized to within-experiment controls, statistical comparisons were conducted using a one-sample t-test with Bonferroni correction.

## RESULTS

### Inconsistent effects of VRAC blockers on cell proliferation in serum-containing cell culture media

Our previous studies using an RNAi-approach established an important role of the LRRC8A protein and LRRC8A-containing VRACs in GBM proliferation (Fidaleo et al., 2026; Rubino et al., 2018). Nevertheless, we and others found that VRAC inhibitors, including the most selective DCPIB, produce inconsistent or no effects on cell proliferation (Fidaleo et al., 2026; Liu and Stauber, 2019). To address these paradoxical results, we comparatively tested five structurally diverse small molecular weight compounds with established effects on VRAC activity. We compared the effects of the best-in-class VRAC inhibitor DCPIB to the recently discovered potent VRAC blocker bromadiolone (Chen et al., 2024), the dihydrochalcone polyphenol phloretin (Fan et al., 2001), and the broad-spectrum chloride channel inhibitors carbenoxolone and DIDS (Benfenati et al., 2009; Stott et al., 2014). Working concentrations of these compounds were selected based on their reported IC_50_ values (see Table 1) and our own work (Abdullaev et al., 2006) and were all expected to produce at least 80% inhibition of VRAC activity.

**Table 1.** Pharmacological VRAC inhibitors and their effective concentrations.

| Pharmacological inhibitor | Reported VRAC IC<sub>50</sub> | Maximal inhibition/concentration | Reference | Maximal water solubility |
| --- | --- | --- | --- | --- |
| Bromadiolone | 1.7 μM | 100% at 10 μM | Chen et al., 2024 | ~25 μM (pH 7.0) |
| Carbenoxolone | 15 μM | >80% at 300 μM | Benfenati et al., 2009 | ~100 mM |
| DCPIB | 4.1 μM | >85% at 10 μM | Decher et al, 2001 | ~50 μM |
| DIDS | 1-20 μM @ MP(+)<br>30-300 μM @ MP(-) | 20-60% at 500 μM (depending on MP) | overview in Okada et al., 2019 | 18 mM |
| Phloretin | 30 μM | >90% at 300 μM | Fan et al., 2001 | 150-500 μM |

**Abbreviations:** DCPIB, 4-[(2-butyl-6,7-dichloro-2-cyclopentyl-2,3-dihydro-1-oxo-1H-inden-5-yl)oxy]butanoic acid; DIDS, 4,4'-diisothiocyanatostilbene-2,2'-disulfonic acid; MP, membrane potential (+) positive or (-).

In the first series of experiments, we utilized an MTT assay to measure changes in relative cell numbers after 72 h incubation with each of the tested compounds. In the fast-proliferating GBM1 cells, the potent VRAC blocker bromadiolone had no effect on cell proliferation (Fig. 1A). All other VRAC inhibitors reduced the MTT signal by 40-70%, with phloretin exerting the strongest inhibition (Fig. 1A). These results were validated using electronic cell counting in a Coulter counter (Fig. 1B) and further qualitatively confirmed through light microscopy observations (Fig. 1C). Curiously, phloretin caused GBM1 to acquire a spindle-like morphology, while DIDS-treated cells grew in distinct cell islands (Fig. 1C).

**Figure 1.**
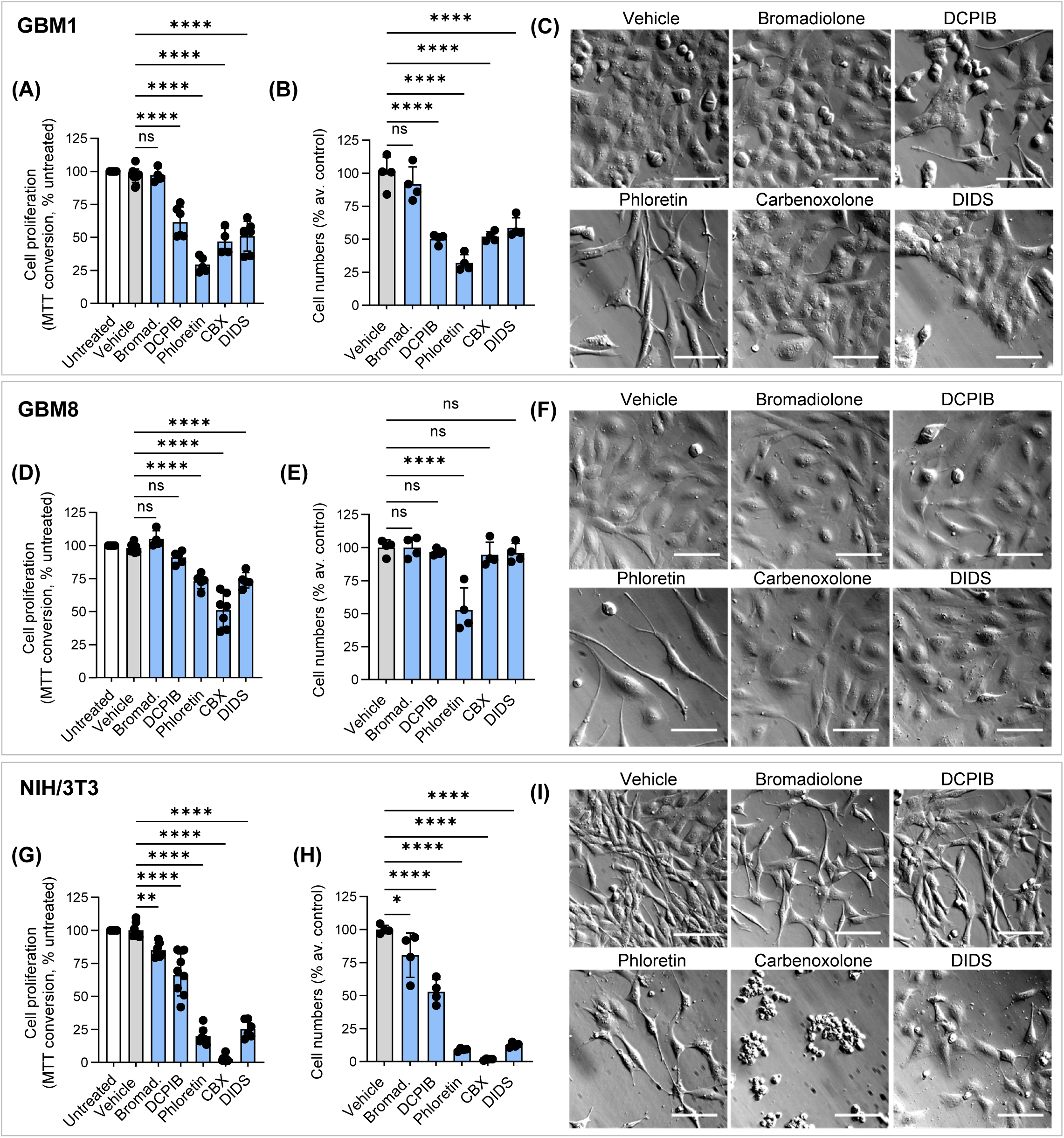
Pharmacological inhibitors of VRAC produce inconsistent effects on cell proliferation in standard cell culture medium. **(A-C)** Effect of VRAC blockers on proliferation of GBM1 cells. Cells were treated for 72 h with 10 µM bromadiolone, 20 µM DCPIB, 200 µM phloretin, 100 µM carbenoxolone (CBX), or 500 µM DIDS. **(A)** Results of MTT cell proliferation assay normalized to an untreated control in the same experiment. Data are mean values ±SD in 5-8 independent experiments. **(B)** Coulter counter counts of cells grown under the same conditions. Mean values of 4 independently treated samples normalized to an average control. **(C)** Representative images of cells treated with VRAC blockers for 72 h. **(D-F)** Effect of VRAC blockers on proliferation of GBM8 cells measured using MTT assay (**D**, n=5-8) or Coulter counter (**E**, n=4). **(F)** Representative images of GBM8 cells captured at 72 h. **(G-I)** Effect of VRAC blockers on proliferation of NIH/3T3 immortalized fibroblasts measured using MTT assay (**G**, n=7-8) or Coulter counter (**H**, n=4). **(I)** Representative images of NIH/3T3 cells captured at 72 h. For all proliferation assays: *p<0.05, **p<0.01; ****p<0.0001, ns, not significant vs. vehicle-treated cells; One-way ANOVA with Dunnett’s multiple comparisons test. Scale bar in all representative images =50 µm.

When the same treatments were tested in the slower growing GBM8 cells, we observed different outcomes. In MTT assays, the most potent inhibitors, bromadiolone and DCPIB, were completely ineffective, phloretin and DIDS inhibited GBM8 growth by ∼25%, while carbenoxolone reduced cell numbers by ∼50% (Fig. 1D). Again, these findings were corroborated using a Coulter counter (Fig. 1E) and light microscopy (Fig. 1F). In GBM8, phloretin produced a spindle-like morphological transformation resembling that seen in GBM1 (Fig. 1F).

We next explored the effects of the same pharmacological agents in a non-malignant NIH/3T3 fibroblast cell line. In immortalized fibroblasts, bromadiolone was largely ineffective (∼15-20% inhibition), DCPIB reduced cell numbers by ∼35-50%, phloretin and DIDS suppressed cell growth by 70-80%, and carbenoxolone killed nearly all the cells (Fig. 1G, H). Light microscopy observations suggested that carbenoxolone was highly toxic to NIH/3T3 cells, causing their rounding and detachment from the substrate (Fig. 1I).

In summary, while some VRAC blockers reduced cell proliferation in serum-containing media, their effects were inconsistent with reported potencies and varied markedly across the three tested cell lines.

### VRAC activity assay indicates that binding to albumin limits the inhibitory potency of VRAC blockers

Next, we tested the efficacy of VRAC blockers in a radiotracer release assay measuring VRAC activity as the efflux of preloaded D-[^3^H]aspartate. Unlike electrophysiology, this approach allows us to assess VRAC function in large populations of intact cells and is compatible with a variety of cell culture conditiuons. Because the majority of VRAC inhibitors are weak lipophilic acids, we speculated that their activity may be limited by binding to albumin in serum-containing media (Ghuman et al., 2005; Valko et al., 2003). Therefore, we performed our experiments in either standard cell culture medium supplemented with 10% FBS, serum-free DMEM, or DMEM supplemented with 4 mg/mL BSA, which approximately corresponds to the albumin content in 10% FBS.

The representative experiments shown in Fig. 2A-C, show that hypoosmotic swelling of GBM1 cells caused a significant release of the preloaded D-[^3^H]aspartate radiotracer, which has been linked to the activation of LRRC8A-containing VRAC (Abdullaev et al., 2006; Chandler et al., 2026; Hyzinski-Garcia et al., 2014). In serum containing medium, the highly potent DCPIB and bromadiolone caused marginal or no inhibition of VRAC activity (Fig. 2A, D). Similarly, carbenoxolone was without effect (Fig. 2D). In comparison, phloretin and DIDS were partially effective and reduced swelling-stimulated release by ∼50-60% (Fig. 2A, D). In striking contrast to results in serum-containing media, in the serum-free DMEM, bromadiolone and DCPIB inhibited VRAC by 82% and 100%, respectively (Fig. 2B, D). Phloretin- and DIDS-induced inhibition in the absence of serum remained approximately the same as in the serum-containing medium (48%, Fig. 2B, D). The addition of albumin to DMEM completely negated the inhibitory actions of DCPIB, bromadiolone, and largely abolished the effect of carbenoxolone (Fig. 2C, D), strongly suggesting that binding to this protein limits the efficacy of these compounds. Conversely, albumin did not modify VRAC inhibition by phloretin or DIDS (Fig. 2C, D).

**Figure 2.**
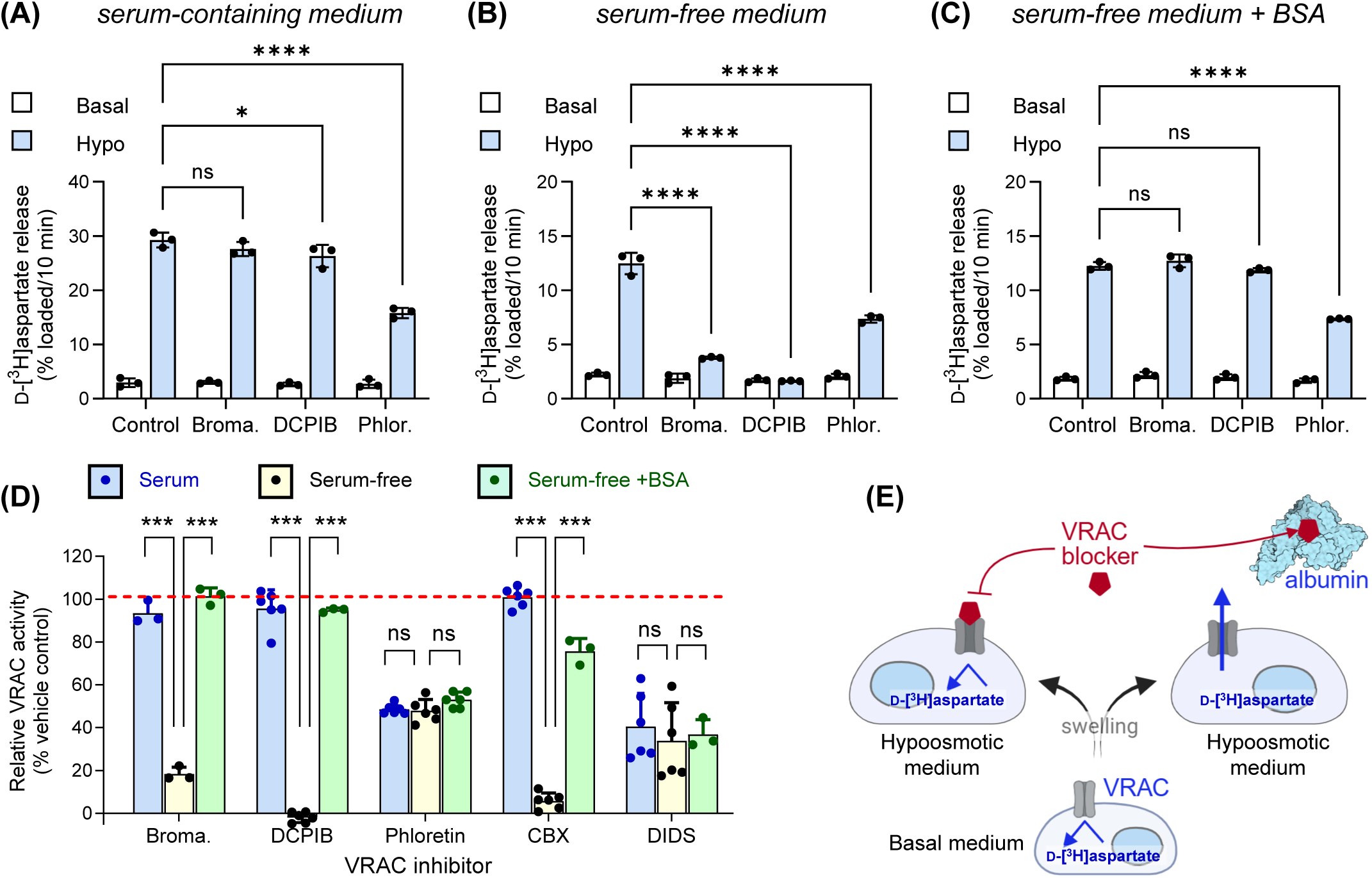
Binding to albumin is a rate-limiting factor for the inhibitory potency of VRAC blockers in cell culture media. **(A-C)** Representative experiments measuring activity of LRRC8A-containing VRACs in the medium containing 10% FBS (**A**), serum-free medium (**B**), or serum-free medium additionally supplemented with 4 mg/mL bovine serum albumin (BSA, **C**), all performed in the same passage of GBM1 cells. VRAC activity was assessed as swelling-activated D-[^3^H]aspartate release in the presence of vehicle 0.1% DMSO (Control), 10 µM bromadiolone (Broma.), 20 µM DCPIB, or 200 µM phloretin (Phlor.). The results are the mean values ±SD of three independent assays performed in quadruplicate. ns, not significant, *p<0.05, ****p<0.0001 vs. hypoosmotic stimulated release in vehicle-treated cells; ANOVA with Tukey correction for multiple comparisons. **(D)** Summary of the effects of bromadiolone (Broma.,10 µM ), DCPIB (20 µM), phloretin (Phlor., 200 µM), carbenoxolone (CBX, 100 µM), or DIDS (500 µM) in serum-containing, serum-free, and serum-free media supplemented with BSA. Data and means ±SD from 3-6 independent experiments. *p<0.05, vs, effect of VRAC blocker in serum-containing and albumin containing medium, one-way ANOVA with Tukey correction for multiple comparison. For clarity, only within-group comparisons are shown. **(E)** Graphical explanation of results shown in (A-D), suggesting that binding to albumin is the limiting factor for efficacy of VRAC blockers.

Overall, these results indicate that binding to albumin is the limiting factor for the efficacy of VRAC blockers in serum-containing growth media (Fig. 2E).

### In serum-free cell culture medium, VRAC blockers potently reduce cell numbers with morphological signs of toxicity

To further examine the effects of VRAC inhibitors without the constraint of binding to serum albumin, we performed additional proliferation assays in GBM cells grown in OptiMEM with added G-5 growth supplement. NIH/3T3 cells did not proliferate in the presence of G5; therefore, we performed fibroblast proliferation assays in low-serum conditions, in OptiMEM medium supplemented with 1% FBS.

In striking contrast with the serum-containing medium, both the MTT assays and Coulter counts found a dramatic reduction in GBM1 and GBM8 cell numbers in the presence of all VRAC blockers except for DIDS (Fig. 3A, B, D, E). Light microscopy observations pointed to significant toxicity of bromadiolone, DCPIB, and carbenoxolone. These compounds caused cell rounding and necrotic cell swelling, although a few shrunken apoptotic cells with membrane blebbing were also observed (Fig. 3C, F). Treatment with phloretin also produced strong toxicity, but in this case some surviving GBM1 and especially GBM8 cells persisted for 72 hours (Fig. 3C, F). DIDS reduced GBM1 numbers by 25% and 56% in MTT and Coulter assays, respectively (Fig. 3A, B). In GBM8 the effects of DIDS were muted: no observable effect in the MTT assay and 23% reduction in Coulter counts (Fig. 3D, E). Consistent with cell counts, DIDS did not cause signs of overt cell death with cell morphology remaining largely similar to serum-free controls (Fig. 3F).

**Figure 3.**
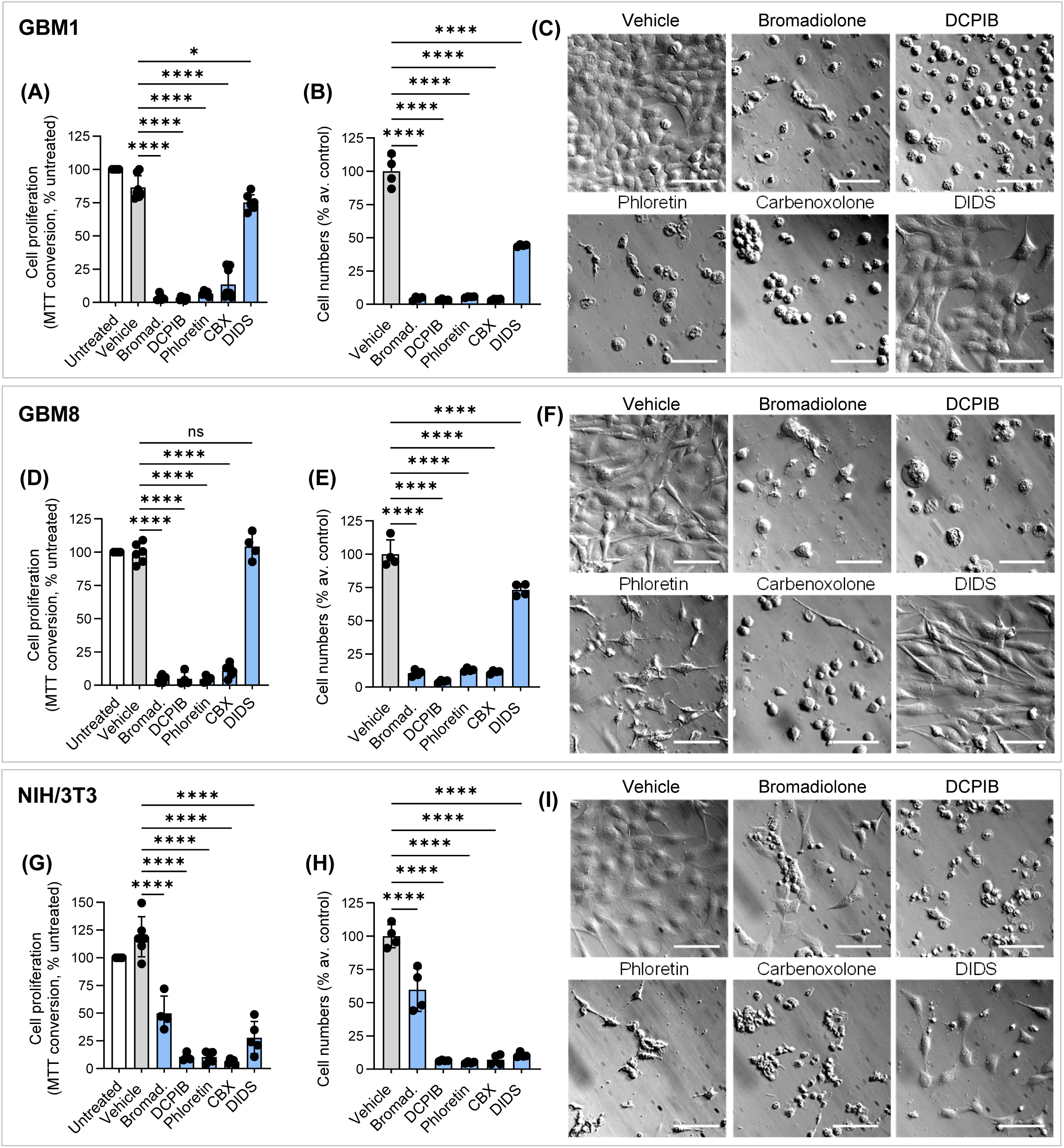
Pharmacological inhibitors of VRAC readily reduce cell proliferation and viability in serum-free cell culture media. **(A-C)** Effect of VRAC blockers on proliferation of GBM1 cells. Cells were treated for 72 h with 10 µM bromadiolone, 20 µM DCPIB, 200 µM phloretin, 100 µM carbenoxolone (CBX), or 500 µM DIDS in OptiMEM supplemented with G5 or 1% FBS (for NIH/3T3). **(A)** Results of MTT cell proliferation assay normalized to an untreated control in the same experiment. Data are mean values ±SD in 5-8 independent experiments. **(B)** Coulter counter counts of cells grown under the same conditions. Mean values of 4 independently treated samples normalized to an average control. **(C)** Representative images of cells treated with VRAC blockers for 72 h. capture cells. **(D-F)** Effect of VRAC blockers on proliferation of GBM8 cells measured using MTT assay (**D**, n=5-8) or Coulter counter (**E**, n=4). (**F)** representative images of GBM8 cells captured at 72 h. **(G-I)** Effect of VRAC blockers on proliferation of NIH/3T3 immortalized fibroblasts measured using MTT assay (**G**, n=4-8) or Coulter counter (**H**, n=4). **(I)** Representative images of NIH/3T3 cells captured at 72 h. For all proliferation assays: *p<0.05, **p<0.01; ****p<0.0001, ns, not significant vs. vehicle-treated cells; One-way ANOVA with Dunnett’s multiple comparisons test. Scale bar in all representative images =50 µm.

In nonmalignant NIH/3T3 cells cultivated in the presence of 1% FBS, all VRAC blockers, including DIDS, dramatically reduced cell numbers (Fig. 3G, H). DCPIB and carbenoxolone caused massive cell rounding (Fig. 3I). Phloretin was also toxic, with signs of necrosis and apoptosis but some cells retaining the spindle-like morphology resembling that seen in the presence of serum (Fig. 3I, compare to Fig. 1I). DIDS left most of the cells intact, but some shrunken apoptotic cells were seen as well (Fig. 3I). Curiously, bromadiolone was the least effective inhibitor in fibroblasts and decreased cell numbers by 50-60% only (Fig. 3G, H). This is likely explained by binding of bromadiolone to albumin in the presence of 1% FBS medium. Light microscopy analysis showed many live cells in the presence of bromadiolone with dying cells exhibiting signs of shrinkage rather than necrotic swelling as in GBM cells (Fig. 3I).

In general, when tested in serum-free or low-serum conditions VRAC blockers, except for DIDS, produced significant toxicity in all three tested cell lines. Partial DIDS toxicity was seen in NIH/3T3 fibroblasts.

### In serum-free media, VRAC blockers deplete intracellular ATP and promote cell death

Since the addition of VRAC blockers in serum-free medium produced overt signs of cytotoxicity, we first quantified cell death levels using an LDH release assay. GBM1 cells were exposed to a panel of VRAC inhibitors for 24 h, and LDH release into the culture medium was normalized to the total LDH content measured in the same wells. As shown in representative images in Fig. 4A, all tested pharmacological agents except DIDS induced pronounced morphological changes in GBM1 cells, similar to those observed after 72 h of treatment (compared to Fig. 3C). These morphological alterations correlated with increased LDH release. At 24 h, we found that 36% of cells lost their viability in the presence of either bromadiolone or DCPIB (Fig. 4B). Phloretin and carbenoxolone increased cell lysis by 16% and 18%, respectively (Fig. 4B). DIDS treatment did not elevate LDH release levels over untreated cells or vehicle-treated controls (Fig. 4B). As a positive control, we used 0.5 µM staurosporine, a well-known inducer of cell death (Bertrand et al., 1994; Jacobsen et al., 1996). Within 24 h, staurosporine caused a 41% increase in LDH release over matching controls (Fig. 4B). Together, these results point to significant acute toxicity of the majority of VRAC blockers, with the cell death levels approaching those seen with the highly toxic staurosporine. This toxicity likely fully accounts for the reduced cell numbers reported at 72 h treatment time in Fig. 3.

**Figure 4.**
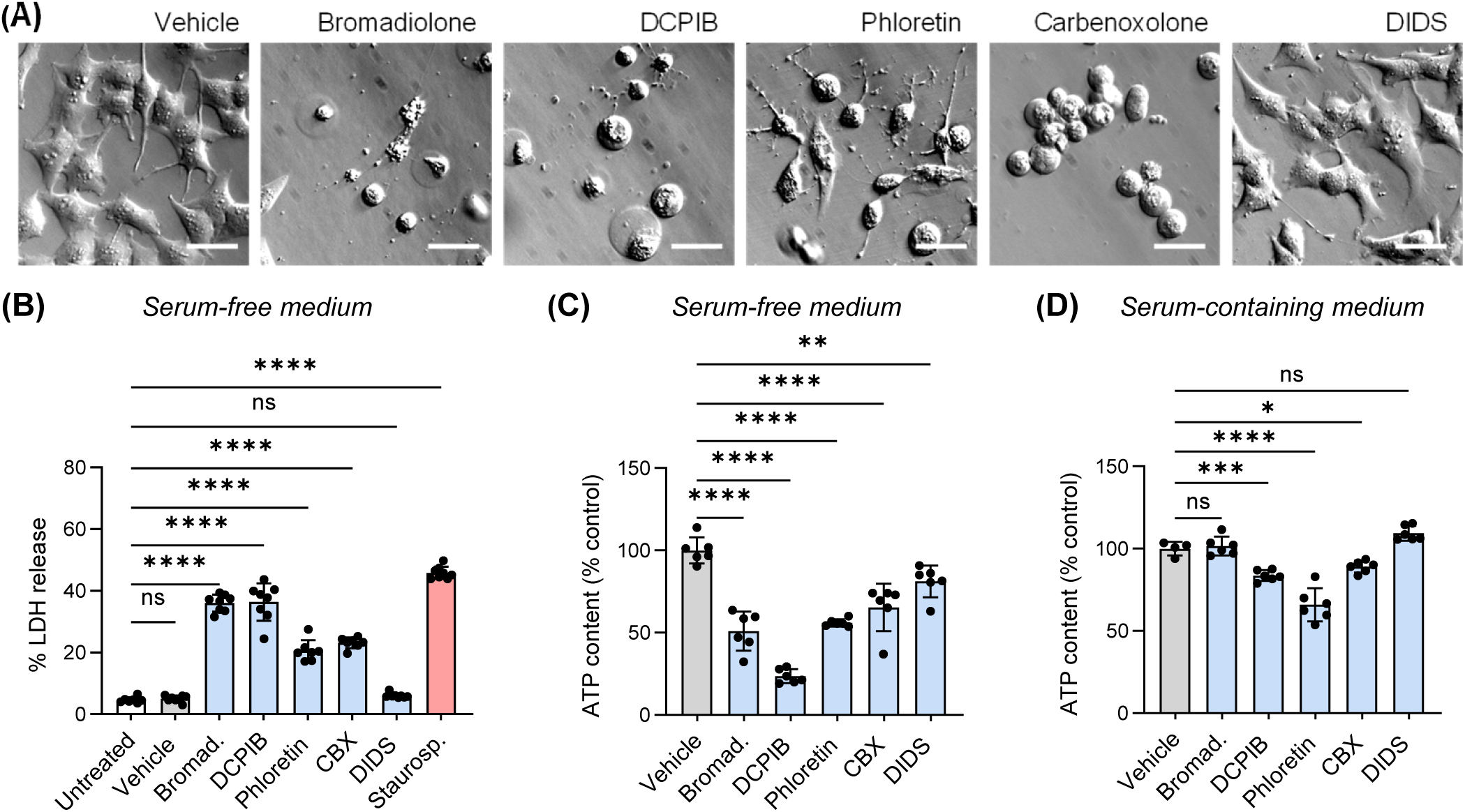
Metabolic inhibition and cell death in GBM1 cells treated with VRAC blockers in serum-free medium. **(A)** Representative images of GBM1 cells treated with VRAC inhibitors in serum-free conditions for 24 h. **(B)** Cytotoxic effect of VRAC blockers in GBM1 cells. Cells were treated with VRAC blockers in serum-free DMEM for 24 h and cell death was assessed as the release of lactate dehydrogenase (LDH), normalized to the total LDH content in each well. 0.5 µM staurosporine (Staurosp.) was used as positive control. (**C)** Effect of VRAC blockers on ATP content of GBM1 cells grown in serum-free medium. Cells were treated for 24 h with 10 µM bromadiolone (Bromad.), 20 µM DCPIB, 200 µM phloretin, 100 µM carbenoxolone (CBX), or 500 µM DIDS. (**D)** Effect of VRAC blockers on ATP content of GBM1 cells treated for 24 h in serum-containing DMEM. For all assays: *p<0.05, **p<0.01; ***p<0.001; ****p<0.0001, ns, not significant vs. vehicle-treated cells; One-way ANOVA with Dunnett’s multiple comparisons test. Scale bar in all representative images =25 µm.

**Figure 5.**
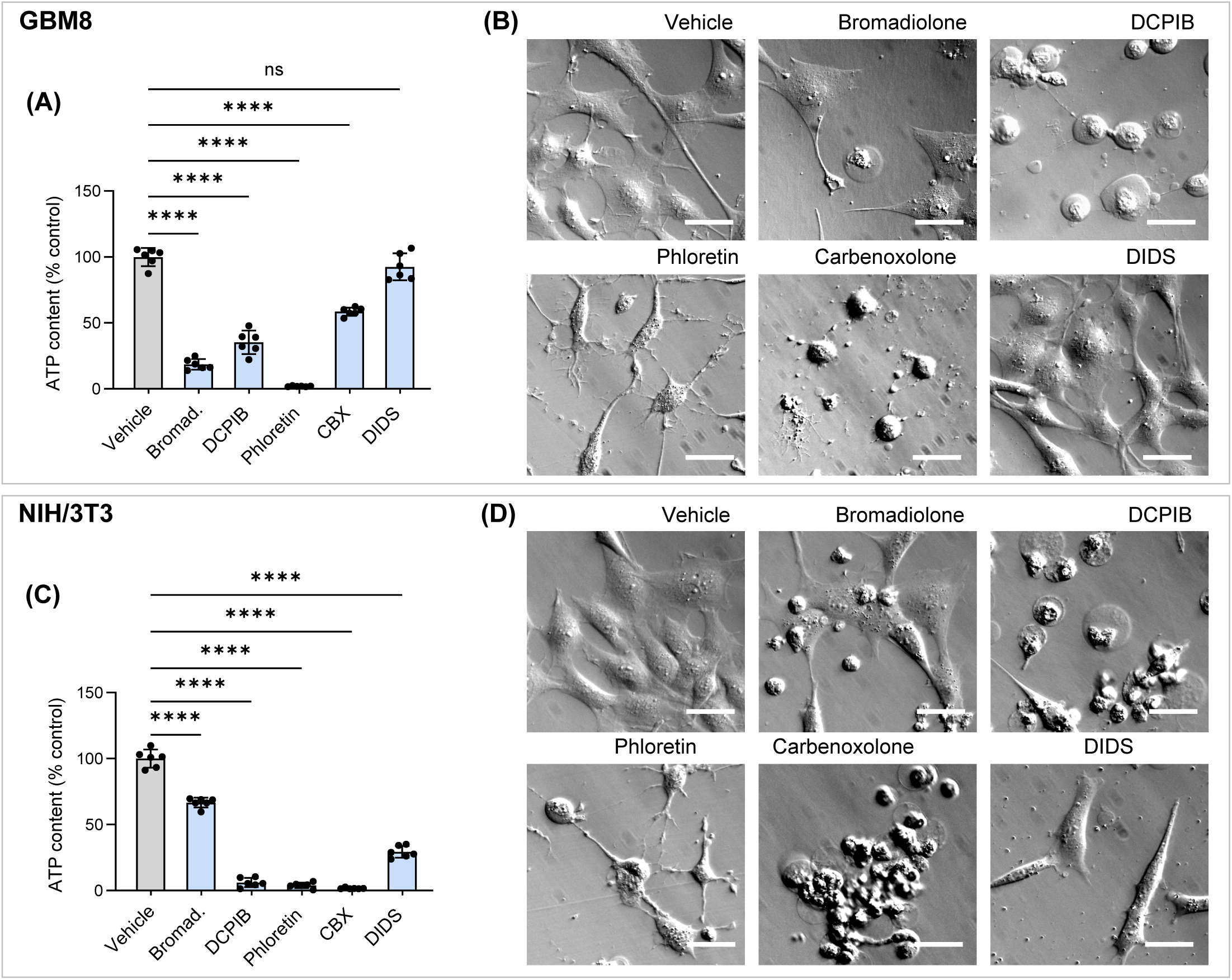
VRAC blockers deplete intracellular ATP and promote cell death in multiple cell types. **(A)** Effect of VRAC blockers on ATP content on GBM8 in serum-free medium. Cells were treated for 24 h with 10 µM bromadiolone (Bromad.), 20 µM DCPIB, 200 µM phloretin, 100 µM carbenoxolone (CBX), or 500 µM DIDS diluted in OptiMEM + G5. **(B)** Representative images of GBM8 cells treated with VRAC inhibitors in serum-free conditions for 24 h. **(C)** Effect of VRAC blockers on ATP content on NIH/3T3 cells in low-serum medium. Cells were treated for 24 h with 10 µM bromadiolone, 20 µM DCPIB, 200 µM phloretin, 100 µM carbenoxolone (CBX), or 500 µM DIDS diluted in OptiMEM + 1% FBS. **D**, Representative images of NIH/3T3 cells treated with VRAC inhibitors in low-serum conditions for 24 h. For all assays: *p<0.05, **p<0.01; ****p<0.0001, ns, not significant vs. vehicle-treated cells; One-way ANOVA with Dunnett’s multiple comparisons test. Scale bar in all representative images =25 µm.

Several studies have reported that VRAC blockers, particularly DCPIB, impair mitochondrial function and inhibit cellular energy metabolism (Afzal et al., 2019). Therefore, we examined whether the VRAC inhibitors used in this study reduce intracellular ATP levels. As an initial control, we tested whether any of the compounds interfered with the luciferase-based ATP detection assay. Under cell-free conditions, the majority of VRAC blockers had no effect on luciferase activity (Supplemental Fig. 1). The only exception was DIDS, which reduced the luciferase signal by approximately 25% after a standard 10-min incubation and by 56% after the extended 2-h incubation (Supplemental Fig. 1), suggesting that the effects of DIDS on ATP production can be overestimated. In serum-free medium, all VRAC blockers produced significant metabolic inhibition in GBM1 cells. After 24 h, the highly cytotoxic bromadiolone and DCPIB reduced intracellular ATP levels by 49% and 76%, respectively (Fig. 4C). Phloretin and carbenoxolone decreased ATP production, by 44% and 35%, respectively (Fig. 4C). DIDS produced a modest effect, lowering intracellular ATP by 19% (Fig. 4C). In contrast, when the same treatments were performed in serum-containing medium, intracellular ATP levels were largely preserved. Phloretin had the highest impact and reduced ATP_i_ by 34% (Fig. 4D). DCPIB and CBX were largely ineffective and reduced ATP content by 16 and 11%, respectively, while bromadiolone and DIDS exerted no effects on ATP_i_ (Fig. 4D). This is likely explained by biding to albumin as demonstrated in VRAC activity assays.

Altogether, the results of LDH release and ATP assays demonstrate that, in serum-free media, VRAC blockers do not merely slow cell proliferation but rather compromise cell viability through disruption of cellular metabolism.

### RNAi-driven downregulation of the essential VRAC subunit LRRC8A inhibits cell proliferation without cytotoxicity

To further determine whether depletion of VRAC induces cell death, we performed additional molecular studies in which VRAC activity was abolished by RNAi-mediated knockdown of the essential VRAC subunit, LRRC8A. Although similar experiments were conducted in our previous work (Fidaleo et al., 2026; Rubino et al., 2018), in the present study GBM1 cells were cultured in serum-free medium supplemented with G-5 to replicate the conditions under which pharmacological VRAC blockers displayed maximal toxicity. We used two siRNA constructs, siLRRC8A_3 (hereafter siA3) and siLRRC8A_5 (siA5), previously validated to efficiently reduce LRRC8A expression and suppress VRAC activity (Fidaleo et al., 2026). As in the prior work, LRRC8A knockdown reduced LRRC8A protein levels by >95% (representative Western blot in Fig. 6A). Consistent with our earlier findings, siA3 and siA5 reduced GBM1 proliferation by 40% and 34%, respectively (Fig. 6B). Importantly, this decrease in cell growth was not accompanied by overt signs of cytotoxicity or morphological alterations (Fig. 6D). As a toxicity control, GBM1 cells were treated with 0.5 µM staurosporine, which caused complete cell death after 72 h (Fig. 6B; representative image in Fig. 6D).

**Figure 6.**
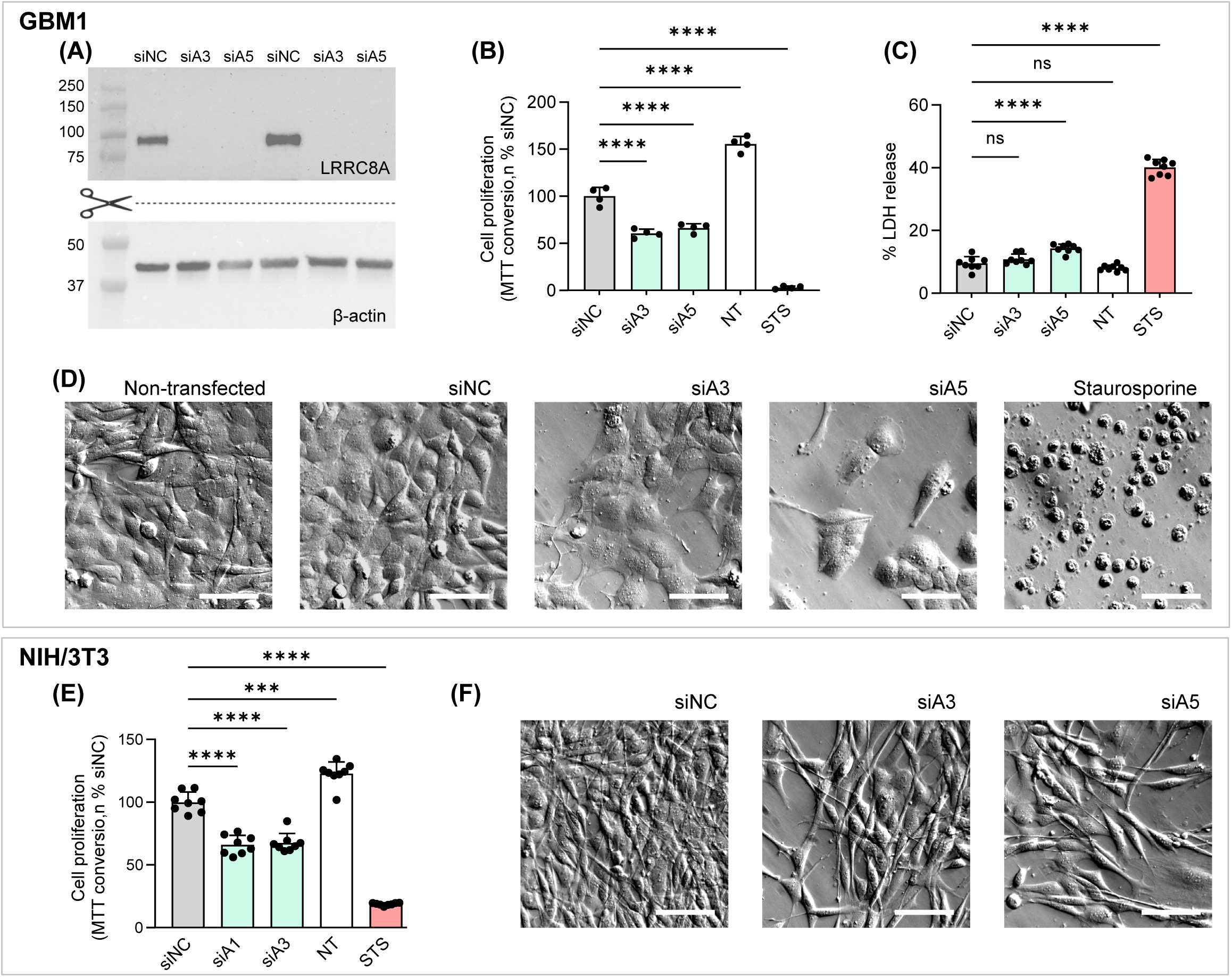
RNAi-mediated knockdown of the essential VRAC subunit LRRC8A inhibits cell proliferation without cytotoxicity. **(A)** Representative western blot image probing downregulation of LRRC8A in GBM1 cells treated with the LRRC8A-specific siRNAs, siA3 and siA5, and compared to the negative control siRNA (siNC). Lower inset shows the same membrane probed for β-actin immunoreactivity. **(B)** Effect of LRRC8A knockdown with the LRRC8A-specific siRNAs siA3 and siA5 on relative proliferation in GBM1 in serum-free medium measured using an MTT assay. 0.5 µM Staurosporine (STS) was used as a control for cell death. Mean values ±SD from four independent transfections. One-way ANOVA with Dunnett’s multiple comparisons test, ****p<0.0001 vs. siNC. **(C)** Minimal cytotoxic effect of LRRC8a knockdown in GBM1 cells. Cells were treated with siNC, siA3 and siA5 in OptiMEM + G5 for 96 h and cell death was assessed as the release of lactate dehydrogenase (LDH), normalized to total LDH content in each well. 0.5 µM Staurosporine was used as positive control for cell death**. (D)** Representative images of GBM1 cells transfected with siRNA for 96h or treated with 0.5 µM staurosporine for 72h. **(E)** Effect of LRRC8A knockdown with the LRRC8A-specific siRNAs siA1(m) and siA3(m) on relative proliferation in NIH/3T3 cells in DMEM +10% FBS measured using an MTT assay. Staurosporine (STS) was used as a control for cell deathl. **(F)** Representative images of NIH/3T3 cells transfected with LRRC8A siRNA for 96h. For all assays: *p<0.05, **p<0.01; ****p<0.0001, ns, not significant vs. vehicle-treated cells; One-way ANOVA with Dunnett’s multiple comparisons test. Scale bar in all representative images =50 µm.

To quantitatively assess whether LRRC8A downregulation causes cytotoxicity, we performed an LDH release assay. Although the siRNA transfection procedure itself caused a slight increase in extracellular LDH, no statistically significant difference was observed between non-transfected and siNC-transfected cells (Fig. 6C). LRRC8A knockdown with siA3 did not produce any additional increase in extracellular LDH, whereas siA5-transfected cells exhibited a modest 4.8% increase in LDH release relative to siNC-treated controls (Fig. 6C). These effects contrasted sharply with the pronounced cell loss and robust LDH release induced by the four VRAC blockers under identical experimental conditions (compare Fig. 6C with Fig. 4B). As a positive control, treatment with staurosporine resulted in 41% LDH release (Fig. 6C).

To determine whether the antiproliferative effect of LRRC8A knockdown also extends to non-malignant cells, we performed MTT proliferation assays in NIH/3T3 fibroblasts. Cells were transfected with the mouse-specific LRRC8A siRNA constructs siA1(m) and siA3(m), which have previously been validated to reduce LRRC8A expression and inhibit VRAC activity (Chandler et al., 2026). Both siA1(m) and siA3(m) reduced cell proliferation by approximately 33% (Fig. 6E). Consistent with our findings in GBM1 cells, this reduction in cell number was not accompanied by obvious signs of cell death, as illustrated by the representative images in Fig. 6F.

Taken together, these findings demonstrate that LRRC8A depletion and consequent loss of VRAC activity significantly suppress cell proliferation, but do not reproduce the pronounced cytotoxic effects observed with pharmacological VRAC blockers.

## DISCUSSION

There are three principal findings in this study. First, molecular biology evidence suggests that, in clear agreement with our prior work, LRRC8A-containing VRACs promote cell proliferation in both malignant glioblastoma and non-malignant NIH/3T3 fibroblast cell line. Second, the majority of VRAC blockers readily bind to albumin, and this interaction explains the discrepancy between their apparent channel affinity in biophysical assays and in cell physiology studies, with the latter typically conducted in serum-containing media. Third, in longitudinal cell physiology experiments, the majority of VRAC blockers exhibit cytotoxicity, which is particularly pronounced under serum-free conditions and is associated with metabolic inhibition. Because the cytotoxic effects of VRAC blockers are not recapitulated in cells with RNAi-mediated depletion of the essential VRAC subunit LRRC8A, the observed cytotoxicity develops independently of VRAC function.

As discussed in the Introduction, the role of VRACs in cell proliferation remains controversial, with conflicting publications both supporting and refuting their importance in this process. Numerous pharmacological studies using putative VRAC inhibitors—including NPPB, DIDS, tamoxifen, and the more selective DCPIB—concluded that inhibition of VRAC activity suppresses proliferation in a variety of cell types [e.g., (He et al., 2012; Shen et al., 2000; Voets et al., 1995; Wondergem et al., 2001)], including glioblastoma cells that are central to the present work (Rouzaire-Dubois et al., 2004). These studies generally attributed the antiproliferative effects of VRAC inhibition to cell-cycle arrest [e.g., (Shen et al., 2000) and reviews (Eggermont et al., 2001; Nilius, 2001)]. However, several more recent publications employing genetic and molecular biology approaches challenged this conclusion, reporting little or no effect of LRRC8A deletion on proliferation in several malignant and non-malignant cell lines (Liu and Stauber, 2019; Widmer et al., 2022; Zhang et al., 2022). Conversely, our present findings, together with previous molecular biological evidence (Fidaleo et al., 2026; Konishi et al., 2019; Kurashima et al., 2021; Lu et al., 2019; Rubino et al., 2018; Xu et al., 2022; Zhang et al., 2018), consistently demonstrate that deletion of LRRC8A reduces proliferation in GBM, other cancers, but also in non-malignant fibroblasts. In contrast to the relatively consistent results obtained with RNAi-mediated LRRC8A knockdown, the effects of pharmacological VRAC inhibitors on cell proliferation were poorly correlated with their reported potencies and varied substantially among cell types. Our findings indicate that these discrepancies arise largely from binding to serum albumin, which reduces the effective concentrations of majority of VRAC inhibitors. Another important factor is nonspecific cytotoxicity that becomes particularly evident under serum/albumin-free conditions. Taken together, our findings support the conclusion that VRACs contribute to sustaining cell proliferation, at least in certain cell types. However, the currently available VRAC blockers are unreliable tools for investigating the role of VRACs in this process.

One important observation of this work is demonstration of limited efficacy and inconsistent effects of potent VRAC blockers DCPIB and bromadiolone on cell growth in serum-containing media. Our VRAC activity assays show that in the presence of 10% serum, these two agents do not block VRAC to any appreciable extent. This lack of efficacy is explained by drug binding to albumin as it can be fully recapitulated by introducing 4 mg/ml of albumin into the VRAC activity assay media. Similar observations have been recently reported by Axel Concepcion’s laboratory who found reduced efficacy of DCPIB and dicumarol in the presence of either serum or albumin [preprint (Granados et al., 2026)]. Perhaps this is not surprising because one of the main functions of albumin is transport of hydrophobic substances such as free fatty acids, thyroid hormones, and others (Fanali et al., 2012). Two primary and several additional drug-binding sites have been identified on the albumin molecule, and their preference for hydrophobic compounds with weak acidic properties has been well characterized (Ghuman et al., 2005). The majority of VRAC blockers, including the recently identified dicumarol and bromadiolone, are hydrophobic weak acids. Notably, the albumin-binding site for warfarin and other 4-hydroxycoumarin-derived anticoagulants such as dicumarol and bromadiolone was extensively characterized at the crystallographic level (Ghuman et al., 2005). Consequently, the development of future therapeutic agents targeting VRAC will need to consider their affinity for albumin, as the extensive plasma protein binding can markedly reduce free drug concentration. Interestingly, in our hands, the VRAC inhibitory activities of phloretin and DIDS were unaffected by the presence of serum or albumin, suggesting that these compounds possess low albumin-binding affinity. This observation may provide structural insight for the rational design of future VRAC inhibitors with reduced plasma protein binding.

The removal of serum from cell proliferation media dramatically increased the inhibitory potency of DCPIB, bromadiolone, and carbenoxolone but also unmasked their significant toxicity. In serum-free media, these agents produced marked signs of cytotoxicity within 24 h and killed the majority of cells in three-day proliferation assays. We found morphological signs of both necrosis (detachment and swelling) and apoptosis (shrinkage and blebbing) and further associated VRAC blocker toxicity with the reduced intracellular ATP levels. Under serum-free conditions all VRAC blockers decrease intracellular ATP levels in a manner that was loosely proportional to the degree of cell death. This is not unexpected as DCPIB and the anticoagulant dicumarol have been shown to inhibit mitochondrial respiration in a VRAC-independent fashion (Afzal et al., 2019; Granados et al., 2026). Due to being weak acids, these compounds can produce mitochondrial uncoupling and reduce oxidative phosphorylation by disrupting the proton gradient through the mechanism ascribed to the well-known mitochondrial uncoupler p-trifluoro-methoxyphenyl hydrazone (FCCP). FCCP is a lipophilic weak acid that shuttles protons across inner mitochondrial membrane (Demine et al., 2019). FCCP, DCPIB, and dicumarol produce mitochondrial depolarization and their mitochondrial effects are blunted in the presence of serum or albumin (Granados et al., 2026). In human and mouse T cells DCPIB and dicumarol strongly increase markers of apoptosis. Among all the VRAC blockers tested by us, DIDS had the smallest effect on ATP levels and did not produce overt signs of cytotoxicity in glioblastoma cells. But even DIDS caused metabolic inhibition and some toxicity in NIH/3T3 fibroblast cell line. These results are in line with recent report on DIDS effect on mitochondrial metabolism and NIH/3T3 cell viability (Das et al., 2025). Overall, these observations raise the question of whether the toxicity associated with VRAC blockers reflects an intrinsic property of this class of agents. Reduction of toxicity certainly should be taken into account while developing novel VRAC-targeting therapies.

In conclusion, our findings suggest that current pharmacological VRAC inhibitors, although valuable in biophysical studies, are poorly suitable for many types of longitudinal cell physiology experiments due to binding to albumin and off-target effects on mitochondrial function and perhaps other cellular processes. Until new less toxic VRAC blockers are found, it is preferable to use molecular biology and molecular genetic strategies targeting LRRC8A when investigating VRAC function.

## AUTHOR CONTRIBUTIONS

AAM conceptualized and supervised this study. MAB, AMA, AR, AMF, HS, and MTK conducted all included experiments. MAB, AMA, AR, and AAM analysed and interpreted experimental results. AAM, MAB, AMA, and AR drafted the manuscript. All authors reviewed, edited, and approved the final version of the manuscript.

## ACKNOWLEDGMENTS

The Authors thank Tejaswi Bannuru for contributions to NIH/3T3 cells experiments and Frances Jourd’heuil and Department of Molecular and Cellular Physiology for providing access to Coulter counter.

