## Supplemental Figure 1 for "Binding to Albumin and Off-Target Toxicity Confound the Use of LRRC8/VRAC Channel Blockers in Cell Physiology Assays"

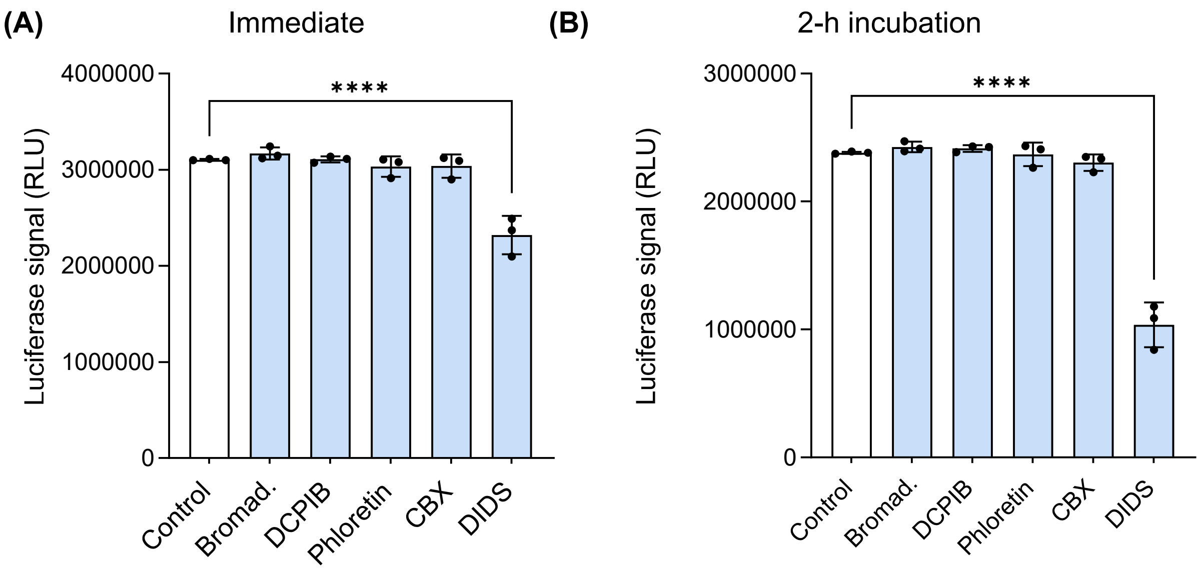


**Effect of VRAC blockers on luciferase activity in the ATPlite assay**

To determine whether VRAC blockers interfere with the ATP detection reagents, control measurements were performed under cell-free conditions. ATP (9 µM) was added to wells containing 10 µM bromadiolone, 20 µM DCPIB, 200 µM phloretin, 100 µM carbenoxolone (CBX), or 500 µM DIDS in OptiMEM + G5 supplement. ATP measurements were performed on the same plate after either a 10-min incubation (**A**) or a 2-h incubation (**B**) in the dark. Data are presented as mean ± SD of replicate measurements from a single experiment. ****$p < 0.0001$ vs. control (one-way ANOVA with Dunnett's multiple-comparisons test).
